# Do l’Hoest’s monkeys show sensitivity to conspecifics’ informational state?

**DOI:** 10.1101/2020.04.29.067686

**Authors:** Lola Rivoal, Guillaume Dezecache, Mélissa Berthet, Audrey Maille

**Author notes:** co-last authorship (Mélissa Berthet, Audrey Maille).

## Abstract

A series of field studies in chimpanzees have shown that the examination of communicative behaviour (e.g. alarm calling) could be particularly fruitful to help uncover mind-reading abilities in non-human animals. In this study, we sought to extend communication-based protocols to one species of Cercopithecids. Specifically, we looked at whether social gazing behaviour of L’Hoest’s monkeys (*Allochrocebus lhoesti*) is sensitive to the informational state of conspecifics using an original experimental design, thereby focusing on their ability to represent conspecifics’ rather than human experimenters’ mental states. We presented a group of 11 zoo-housed monkeys with a box that contained either an appetitive (mealworms), aversive (raptor stuffed toy) or neutral stimulus (wood bark chips). The discoverer (the first individual that inspected the content of the box) emitted more social gazes in the aversive condition than in the neutral and appetitive conditions. Besides, social gazing was related to the informational state of the conspecifics in the appetitive and neutral conditions, with more social gazes directed towards informed individuals (who had inspected the box) rather than uninformed ones (who had not inspected the box yet). Additional examination of the behaviour of the discoverers revealed that they were likely addressing social gazes to individuals who were in close proximity to them, suggesting that they preferentially gaze at informed conspecifics because the latter remained in proximity. Our study calls for a more widespread assessment of mind-reading capacities in primates, to further reveal the evolutionary history of traits that were thought to be uniquely human until fairly recently.

## Introduction

Humans routinely communicate using signals that are tuned to meet others’ epistemic states (i.e., what others know and believe, Sperber and Wilson 1987). This capacity requires two cognitive abilities: (i) the ability to notice and keep track of what others have perceived or are currently able to perceive; (ii) the ability to provide others with information they lack.

Studies suggest that adjustment to the epistemic states of others is not unique to humans. Great apes indeed appear to be sensitive to others’ false beliefs (Krupenye et al. 2016; Kano et al. 2019), although evidence remains discussed (see Heyes 2017; Kano et al. 2017; Krupenye et al. 2017 for discussion). Less controversial are the chimpanzees’ abilities to adjust their behaviour to the informational state of conspecifics when competing for food (Hare et al. 2001; Kaminski et al. 2008). Chimpanzees also modify their alarm calling behaviour to what their conspecifics have seemingly been able to see or hear (Crockford et al. 2012, 2017). Beyond great apes, sensitivity to others’ informational state in primates is widely debated (Heyes 2015; Martin and Santos 2016; Meunier 2017; Krupenye and Call 2019).

Studies have shown that several monkey species are able to track others’ visual perspective, i.e., what others are currently able to perceive (*Macaca fascicularis*, Overduin-de Vries et al. 2014; *Sapajus apella*, Anderson et al. 1995; Hare et al. 2003), and to adjust their communicative behaviour accordingly (*Macaca tonkeana*, Canteloup et al. 2015, 2016; *Cercocebus torquatus*, Maille et al. 2012; *Papio anubis*, Bourjade et al. 2014). Further research has additionally shown that one species of Cercopithecids (*Macaca mulatta*, Marticorena et al. 2011; Drayton and Santos 2017), and one species of Cebids (*Cebus appella*, Kuroshima et al. 2002) are able to track others’ knowledge, i.e. what others have been able to perceive.

Several experimental approaches have been designed to uncover mind-reading capacities in non-human animals. One major methodological choice, particularly suitable for captive animals, has been to explore social cognition via animals’ interactions with human caregivers or experimenters (e.g., Maille et al. 2012). Another has been to trigger competition between same-species individuals (e.g., Hare et al. 2001), relying on the assumption that competitive interactions motivate the subjects to process others’ mental states to behave deceptively. A more recent approach is to confront subjects with situations mimicking ecologically-relevant events that can trigger warning behaviour, such as the discovery of a threat (e.g., Crockford et al. 2012, 2015, 2017; Schel et al. 2013). Such methodological choice has been particularly potent for it relied on relevant ecological pressures, and in response to which communication, collective defence and overall cooperation are expected (Caro 2005).

We used this approach to assess whether L’Hoest’s monkeys (*Allochrocebus lhoesti*) adjust their social behaviour to conspecifics’ informational state, i.e., what conspecifics have been able to perceive. L’Hoest’s monkeys are African forest monkeys (Mittermeier et al. 2013) that are mostly terrestrial and live in small social groups composed of one adult male and 10 to 17 females and their offspring (Kingdon 2015). We confronted a family group of captive L’Hoest’s monkeys with ecologically-relevant stimuli aimed at eliciting communication. For this, we presented the group with an appetitive (palatable food), neutral (wood bark chips) or aversive (a stuffed raptor toy) stimulus, only visible from two openings on the front of a screen. We focused our analyses on social gazes (i.e. the action of looking towards a conspecific, as opposed to non-social gazing directed towards non-living objects) because this behaviour is prominent in a variety of primate species (Tomasello et al. 1998; Emery 2000) and because vocalizations could not be reliably recorded in our zoo setting. We investigated the extent to which the social gazing behaviour of the first animal checking the stimulus (the discoverer) was influenced by the dynamic epistemic states of its conspecifics (whether they had or had not checked the box themselves at the time of the gaze).

Considering the design of our experiment, four hypotheses could be put forward to account for a modulation of social gazing by the informational state of the target conspecific. First, the “proximity” hypothesis (the less cognitively demanding of all four hypotheses) supposes that the discoverer would preferentially gaze at the closest individual(s), which happen to have checked the content of the box because they were close to it. To control for this hypothesis, we assessed whether the discoverer preferentially gazed at the closest individuals. Second, the “saliency” hypothesis supposes that the discoverer would preferentially gaze at informed conspecifics because they would exhibit more conspicuous behaviours than uninformed individuals. While we expect uninformed individuals to perform a baseline behaviour, informed individuals may try to reach the stimulus or emit food-related vocalizations if presented with the appetitive stimulus (while we acknowledge that they may also adopt a cryptic, deceptive behaviour to avoid food competition), or mob, flee away from the testing apparatus or emit alarm calls if presented with the aversive stimulus. This hypothesis can be discarded if the number of social gazes addressed to conspecifics is a function of their epistemic state in the neutral condition, during which we do not expect informed individuals to perform such conspicuous behaviours. Third, the “behaviour reading” hypothesis states that the discoverer acts as if it could process the informative state of conspecifics, but merely reads out of their behaviour, which may look odd. This hypothesis could be discarded if the number of social gazes received by individuals differs with their epistemic state in the neutral condition. Finally, the “mind-reading hypothesis” states that L’Hoest’s monkeys are capable of tracking what others have been able to perceive. Being cognitively more demanding, this hypothesis would be put forward if and only if the three alternative hypotheses can be discarded.

## Materials and methods

### Animals and study site

This study was conducted on a captive group of 11 L’Hoest’s monkeys at La Ménagerie, le zoo du Jardin des Plantes (Museum National d’Histoire Naturelle, Paris). The study group was housed in an enclosure composed of an indoor room connected to 3 outdoor rooms (total size: 27.75 m × 2.50 m × 5.50 m), exposed to visitors every day from 9:00 am to 6:00 pm. The group consisted of one adult male, two related adult females and their offspring, a social structure which resembles the harem structure of L’Hoest’s monkeys in the wild (Kingdon and Largen 1997). Three of the 11 individuals (one female and her offspring) were absent during the seventh first trials of the study for temporary medical isolation (**Table 1**). This study was performed in accordance with institutional ethical guidelines

**Table 1.** Raw data for adult female Hestia and her offspring. Nature of the stimulus for each trial, identity of the discoverer, duration of each trial, total number of gazes that were given during the trial and details about the receivers for these gazes. Abbreviations and codes: Disc. = discoverer; Dur. = duration of the trial (s); Time = time of the trial when the receiver turned informed; Bef. = number of gazes received before turning informed; Aft. = number of gazes received after turning informed; ♀ = female; ♂ = male

| Day | Trial | Stimulus | Disc. | Dur. (s) | Total gazes | Receivers (lineage female Hestia) |  |  |  |  |  |  |  |  |  |  |  |  |  |  |
| --- | --- | --- | --- | --- | --- | --- | --- | --- | --- | --- | --- | --- | --- | --- | --- | --- | --- | --- | --- | --- |
|  |  |  |  |  |  | KIGALI ♂ |  |  | TALA ♀ |  |  | HESTIA ♀ |  |  | KOUMI ♀ |  |  | MINUS ♀ |  |  |
|  |  |  |  |  |  | Time | Bef. | Aft. | Time | Bef. | Aft. | Time | Bef. | Aft. | Time | Bef. | Aft. | Time | Bef. | Aft. |
| 1 | 1 | neutral | Hakan | 543 | 21 | 170 | 2 | 2 | 29 | 0 | 2 | AWAY FROM STUDY |  |  |  |  |  |  |  |  |
| 1 | 2 | appetitive | Hakan | 583 | 34 | 38 | 1 | 7 | 141 | 0 | 2 |  |  |  |  |  |  |  |  |  |
| 2 | 3 | aversive | Hakan | 597 | 75 | - | 15 | - | - | 17 | - |  |  |  |  |  |  |  |  |  |
| 2 | 4 | neutral | - | - | - | Deleted Trial : No discoverer for this trial |  |  |  |  |  |  |  |  |  |  |  |  |  |  |
| 3 | 5 | aversive | Hakan | 564 | 34 | - | 6 | - | - | 3 | - | AWAY FROM STUDY |  |  |  |  |  |  |  |  |
| 3 | 6 | aversive | - | - | - | Deleted Trial : No discoverer for this trial |  |  |  |  |  |  |  |  |  |  |  |  |  |  |
| 4 | 7 | appetitive | Kigali | 543 | 26 |  |  |  | 70 | 1 | 2 | AWAY FROM STUDY |  |  |  |  |  |  |  |  |
| 4 | 8 | appetitive | Kigali | 470 | 20 |  |  |  | - | 3 | - | - | 0 | - | - | 0 | - | - |  | - |
| 5 | 9 | neutral | Hakan | 582 | 34 | - | 6 | - | 38 | 0 | 17 | - | 1 | - | 116 | 2 | 4 | - |  | - |
| 5 | 10 | aversive | Kigali | 587 | 32 |  |  |  | 34 | 0 | 5 | - | 0 | - | - | 11 | - | - | 3 | - |
| 6 | 11 | aversive | Kigali | 319 | 38 |  |  |  | 45 | 0 | 6 | - | 0 | - | 161 | 0 | 1 | - | 1 | - |
| 6 | 12 | appetitive | Hakan | 579 | 34 | - | 1 | - | 270 | 1 | 7 | - | 2 | - | - | 0 | - | - | 2 | - |
| 7 | 13 | appetitive | - | - | - | Deleted Trial : Wrong identification of the discoverer for this trial |  |  |  |  |  |  |  |  |  |  |  |  |  |  |
| 7 | 14 | aversive | Kigali | 587 | 48 |  |  |  | - | 9 | - | - | 6 | - | - | 0 | - | - | 1 | - |
| 8 | 15 | neutral | Hakan | 585 | 21 | 19 | 0 | 2 | - | 0 | - | 258 | 7 | 2 | - | 1 | - | - | 2 | - |
| 8 | 16 | neutral | Kigali | 594 | 16 |  |  |  | 390 | 10 | 0 | - | 0 | - | - | 0 | - | - | 0 | - |
| 9 | 17 | appetitive | Hakan | 587 | 27 | 169 | 0 | 4 | 75 | 2 | 2 | - | 2 | - | 547 | 2 | 2 | - | 1 | - |
| 9 | 18 | neutral | Kigali | 586 | 14 |  |  |  | - | 0 | - | 488 | 7 | 0 | - | 1 | - | - | 0 | - |

### Data collection

Data collection was performed from May 2^nd^ to May 16^th^, 2018. The screen was a cardboard crate (38.5 cm × 29.5 cm × 20.0 cm) with two openings on it (8 cm × 6cm), located 17 cm apart from each other, at 38 cm high and at 5 cm from the left and right edges. The screen was placed outside of the enclosure, against one of the mesh fences, so that the monkeys could not reach the screen nor see what was behind it without looking through one of the openings (Figure 1). The screen remained there throughout the study. Before each stimulus presentation, a caregiver hid one of the stimuli in a round box (diameter = 19.8 cm, height = 17.1 cm) and placed the box behind the screen, allowing the monkeys to check its content from the screen openings. LR recorded and commented each stimulus presentation through the enclosure glass panel opposite to the mesh fence using a Panasonic DMC-FZ300-EF video-camera during up to 600 seconds (trials ranging from 347.42 to 600.00 seconds, mean ± SD = 571.18 ± 69.36 seconds).

**Fig. 1.**
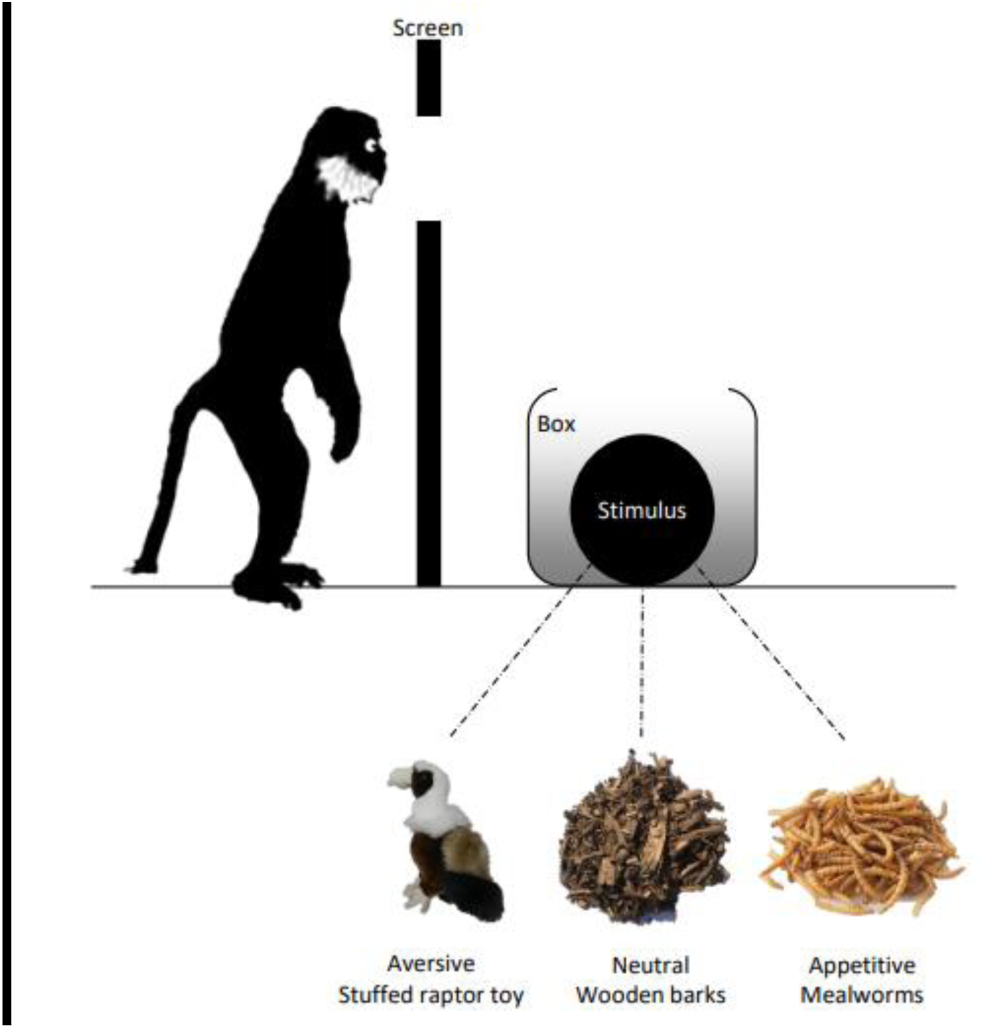
Experimental design. For each trial, a box containing either an aversive (stuffed raptor toy), neutral (wooden barks) or appetitive (mealworms) stimulus was placed behind a screen on which two openings had been cut. The monkeys could inspect the content of the box only by looking through one of the openings.

The box could contain either an appetitive (mealworms, a palatable food item), neutral (mulsh, bark chips litter used in the enclosure) or aversive (a raptor stuffed toy) stimulus (**Figure 1**). The raptor toy was considered as aversive since it elicited a strong negative reaction similar to that of other primate species when encountering a predator in the wild (e.g., prolonged stares) when it was presented to the whole group during a pilot trial performed in a different location.

The experiment was spread over 9 testing days, during which two stimulus presentations were conducted in a row, separated by a 5-minute pause. Each stimulus was presented 6 times in a pseudo-randomized order, so that every possible stimulus combination within a testing day was presented once (**Tables 1, 2**). Note that, for the appetitive condition, the mealworms were distributed to the monkeys at the end of the trials for consumption.

**Table 2.** Raw data for dominant male Hakan, adult female Honey and her offspring. Nature of the stimulus for each trial, identity of the discoverer, duration of each trial, total number of gazes that were given during the trial and details about the receivers for these gazes. Abbreviations and codes: Disc. = discoverer; Dur. = duration of the trial (s); Time = time of the trial when the receiver turned informed; Bef. = number of gazes received before turning informed; Aft. = number of gazes received after turning informed; ♀ = female; ♂ = male

| Day | Trial | Stimulus | Disc. | Dur. (s) | Total gazes | Receivers (reproductive male * + lineage female Honey) |  |  |  |  |  |  |  |  |  |  |  |  |  |  |  |  |  |
| --- | --- | --- | --- | --- | --- | --- | --- | --- | --- | --- | --- | --- | --- | --- | --- | --- | --- | --- | --- | --- | --- | --- | --- |
|  |  |  |  |  |  | HAKAN ♂ * |  |  | HONEY ♀ |  |  | SHELLY ♀ |  |  | BAKA ♂ |  |  | MIYA ♀ |  |  | CORTEX |  |  |
|  |  |  |  |  |  | Time | Bef. | Aft. | Time | Bef. | Aft. | Time | Bef. | Aft. | Time | Bef. | Aft. | Time | Bef. | Aft. | Time | Bef. | Aft. |
| 1 | 1 | neutral | Hakan | 543 | 21 |  |  |  | - | 1 | - | - | 2 | - | 299 | 2 | 9 | - | 1 | - | - | 0 | - |
| 1 | 2 | appetitive | Hakan | 583 | 34 |  |  |  | 139 | 4 | 10 | - | 0 | - | - | 0 | - | - | 5 | - | - | 5 | - |
| 2 | 3 | aversive | Hakan | 597 | 75 |  |  |  | 168 | 3 | 3 | - | 3 | - | - | 16 | - | - | 17 | - | - | 1 | - |
| 2 | 4 | neutral | - | - | - | Deleted Trial : No discoverer for this trial |  |  |  |  |  |  |  |  |  |  |  |  |  |  |  |  |  |
| 3 | 5 | aversive | Hakan | 564 | 34 |  |  |  | - | 6 | - | 42 | 0 | 3 | - | 3 | - | - | 12 | - | - | 1 | - |
| 3 | 6 | aversive | - | - | - | Deleted Trial : No discoverer for this trial |  |  |  |  |  |  |  |  |  |  |  |  |  |  |  |  |  |
| 4 | 7 | appetitive | Kigali | 543 | 26 | 36 | 0 | 10 | - | 4 | - | - | 0 | - | - | 0 | 0 | 200 | 4 | 3 | - | 2 | - |
| 4 | 8 | appetitive | Kigali | 470 | 20 | - | 0 | - | - | 5 | - | - | 0 | - | 71 | 2 | 9 | - | 3 | - | - | 0 | - |
| 5 | 9 | neutral | Hakan | 582 | 34 |  |  |  | - | 1 | - | - | 0 | - | 49 | 1 | 2 | - | 0 | - | - | 0 | - |
| 5 | 10 | aversive | Kigali | 587 | 32 | 15 | 0 | 1 | - | 0 | - | - | 4 | - | - | 4 | - | - | 4 | - | - | 0 | - |
| 6 | 11 | aversive | Kigali | 319 | 38 | 31 | 0 | 17 | - | 0 | - | - | 0 | - | 58 | 0 | 2 | 111 | 0 | 11 | - | 0 | - |
| 6 | 12 | appetitive | Hakan | 579 | 34 |  |  |  | - | 4 | - | - | 7 | - | 364 | 4 | 2 | - | 4 | - | - | 0 | - |
| 7 | 13 | appetitive | - | - | - | Deleted Trial : Wrong identification of the discoverer for this trial |  |  |  |  |  |  |  |  |  |  |  |  |  |  |  |  |  |
| 7 | 14 | aversive | Kigali | 587 | 48 | - | 26 | 0 | - | 1 | - | - | 1 | - | 276 | 2 | 1 | - | 0 | - | - | 0 | - |
| 8 | 15 | neutral | Hakan | 585 | 21 |  |  |  | - | 5 | - | - | 0 | - | 40 | 0 | 1 | 37 | 0 | 1 | - | 0 | - |
| 8 | 16 | neutral | Kigali | 594 | 16 | 13 | 0 | 3 | - | 0 | - | - | 0 | - | 102 | 1 | 2 | - | 0 | - | - | 0 | - |
| 9 | 17 | appetitive | Hakan | 587 | 27 |  |  |  | - | 0 | - | - | 6 | - | 545 | 4 | 0 | - | 2 | - | - | 0 | - |
| 9 | 18 | neutral | Kigali | 586 | 14 | - | 0 | - | - | 0 | - | - | 1 | - | - | 4 | - | - | 0 | - | - | 0 | - |

### Video analysis

Videos were transcribed by LR using BORIS (version 6.2.4) [18]. Trials in which no monkey checked the content of the box (N=2) or for which the video was of poor-quality (N=1) were excluded from the dataset. The analysis was thus performed on 15 out of 18 trials (5 per stimulus type) (**Tables 1, 2**).

We considered that an individual checked the content of the box if the distance between the eyes of this individual and the closest screen opening was shorter than the individual’s head diameter. Individuals were considered as uninformed as long as they had not checked the content of the box. They were considered as informed from the time they first checked the content of the box until the end of the trial. We acknowledge that uninformed individuals probably collected cues about the content of the box from behaviour and vocalizations of informed individuals, and thus, were not completely uninformed. However, they are still to be considered less informed than individuals that did check the box by themselves. It must be noted that 5 (out of 42) checking events were removed from the dataset because the checker could not be identified. We assume that the risk that omission of those events impacted our data on the informational state of the individuals is negligible, since informed individuals often checked the content of the box several times per trial.

The behaviour of the first individual to check the content of the box (hereafter, the discoverer) was transcribed from its first check to the end of the trial (N = 11 trials) or until this individual became not visible (N = 4 trials). For each trial, one observer collected each social gaze given by the discoverer (social gaze defined as the head of the focal individual orienting towards a conspecific) and the identity of the individual that received the gaze (as judged by the head orientation of the discoverer).

Intra-observer reliability was measured based on repeated transcription of 10% of the videos by the same observer over three days. The Fleiss’ Kappa revealed an almost perfect agreement (>0.81).

We later coded whether social gazes were addressed to the closest individual to the discoverer. This was evaluated in a post-hoc fashion, using the video images only, as this was not verbally coded by the observer during the experiment. We were only able to assess spatial proximity in videos where the discoverer was near the centre of the camera view range (∼54% of the social gazes in appetitive and neutral trials; **Supplementary material**).

### Statistical analysis

We analysed how informational state and stimulus type affected the number of gazes that individuals received. We fitted three generalized-linear mixed models for Poisson error structure with, as dependent variable, the number of social gazes received by each individual at each step of its knowledge (i.e. uninformed or informed) for each trial (e.g. the number of social gazes received by individual A in trial 1, while uninformed). In the first GLMM (full model), the fixed factor was the interaction between the informational state of the receiver (uninformed or informed) and the stimulus (aversive, neutral or appetitive). The duration of each informational state (i.e. the duration during which the receiver was either informed or uninformed) was controlled for by including an offset term. Random factors were receivers’ identity (nested in discoverer’s identity) and trial number. We are aware that it would have been important to include trial number as fixed factor to control for habituation, but our dataset prevented us from running more complex analyses: models do not converge anymore if the trial number is a fixed factor.

We compared the full model against a null model (i.e. without fixed factors) and reduced model (i.e. without the interaction between the fixed factors) using likelihood ratio tests (LRT). Post-hoc tests (pairwise t-tests) were conducted with Tukey correction. The fit of the models was evaluated by the proportion of variance explained (the marginal coefficient of determination R2m, i.e. the variance accounted for by fixed factors, and the conditional coefficient of determination R2c, i.e. the variance accounted for by both fixed and random factors). Statistical analyses were carried out using R version 3.4.1 (R Core Team 2019), and the lme4 (Bates et al. 2014) and emmeans (Russell 2018) packages for the GLMM and post-hoc tests respectively. Significance level was set at 0.05. The data and scripts are available as electronic supplementary material.

To evaluate the likelihood that the discoverer preferentially addresses social gazes at the closest individuals, we used binomial tests with varying probability to gaze at this individual among other group members. Since we could not be sure how many individuals were in the vicinity of the discoverer (i.e. gaze behaviour was coded from video footages in which the video range is focused on the discoverer), we estimated the probability that the closest individual is gazed as 1/n, with n ranging from 2 to 10 (total number of conspecifics that could be looked at from the discoverer’s point of view).

## Results

For all of the 15 trials, the discoverer was one of the two highest ranking males of the group (either the reproducing male or the oldest male offspring). Each of these two males was the discoverer for 2 or 3 trials out of 5 trials displaying the same stimulus type. The number of individuals becoming informed during a trial varied from 0 to 5. The number of informed individuals at the end of a trial were 2.8 ± 1.10 (mean ± SE) individuals in the neutral condition, 2.6 ± 1.14 (mean ± SE) in the appetitive condition and 2 ± 1.73 (mean ± SE) in the aversive condition. The number of gazes that each individual received during the 15 trials, depending on its informational state, is presented in **Tables 1 and 2**.

The number of received gazes was higher when the stimulus was aversive than when it was neutral (*t*-test: z ratio = 4.622, p < 0.001) or appetitive (*t*-test: z ratio = 2.830, p = 0.013), regardless of the informational state (Figure 2). There was no difference in the number of gazes received when the stimulus was neutral or appetitive (*t*-test: z ratio = −1.930, p = 0.130).

**Fig. 2.**
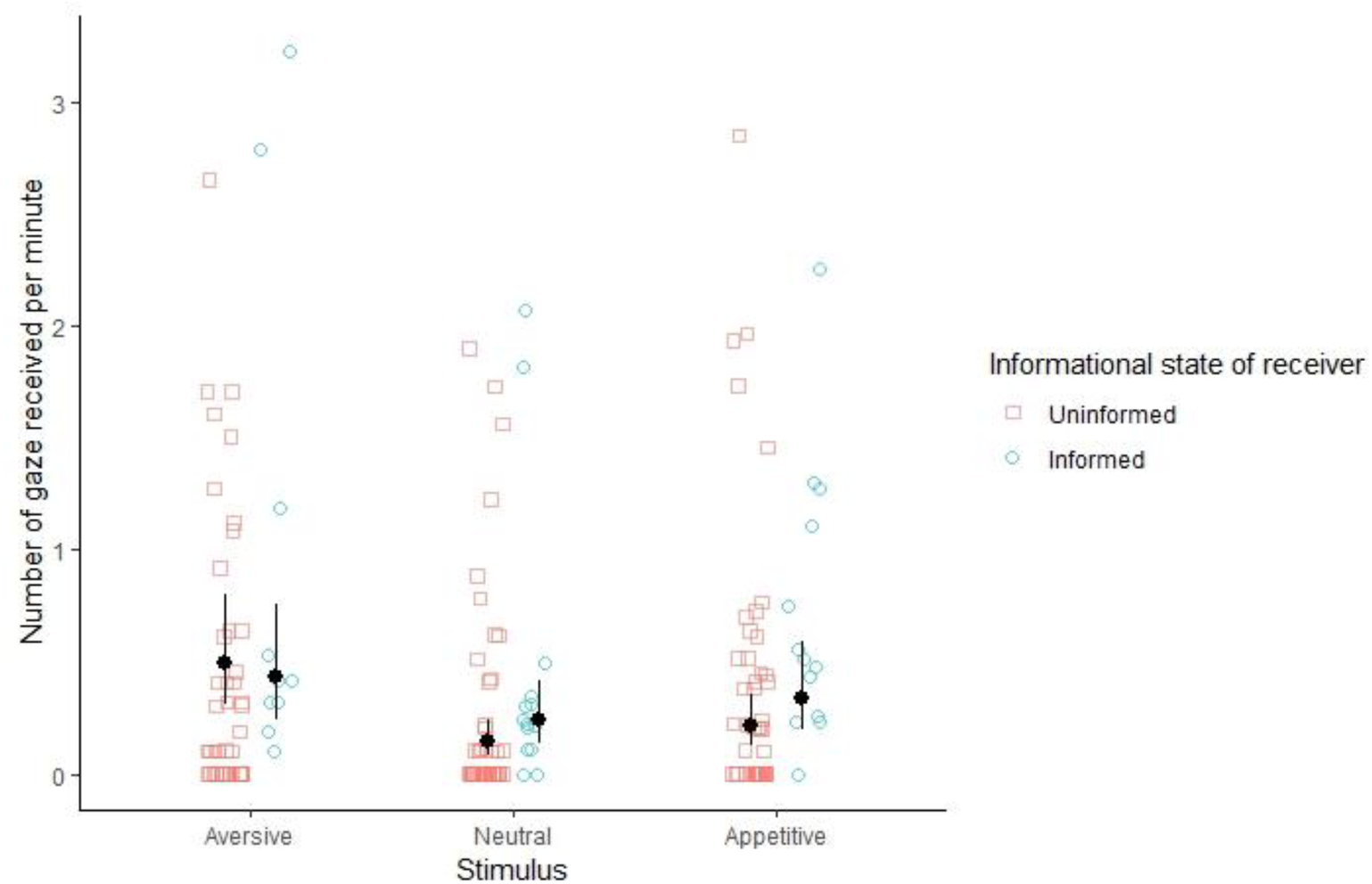
Number of social gazes received per minute as a function of stimulus type and receiver’s informational state. The figure shows raw data (coloured points), model’s estimated marginal means (black dots) and associated 95% confidence intervals of estimates for each condition (black lines).

The interaction of stimulus type and informational state of receiver had an effect on the number of received gazes (i.e. number of gazes addressed to the receiver by the discoverer): the full model (R2m: 0.038; R2c: 0.131) was indeed significantly different from the null (LRT: X2(5) = 26.686, p < 0.001) and the reduced (LRT: X2(2) = 6.971, p = 0.031) models. Informed conspecifics received more gazes than uninformed conspecifics when the stimulus was appetitive (*t*-test: z ratio = −2.436, p = 0.015) or neutral (*t*-test: z ratio = −2.432, p = 0.015), but not when it was aversive (*t*-test: z ratio = 0.757, p = 0.449; Figure 1).

Analyses of 135 gazes coded in the appetitive and neutral conditions revealed that 71 gazes (53 %) were directed towards the individual who was the closest to the discoverer. The closest individual was preferentially gazed at if there were at least three individuals around the discoverer (Binomial test: if probability of success = 1/2, p = 0.60; if probability of success = 1/3, p < 0.001).

## Discussion

When the zoo-housed group of L’Hoest’s monkeys (*Allochrocebus lhoesti*) was presented with a box containing a stimulus in a social context, the adult male discoverers emitted more social gazes when facing an aversive stimulus than appetitive or neutral stimuli. They also gazed more towards informed conspecifics (i.e., conspecifics that had also checked the content of the box) than towards uninformed ones when the stimulus was either appetitive or neutral. Finally, discoverers were more likely to gaze at the closest individual if there were at least three individuals around them.

L’Hoest’s monkeys thus seemed to show sensitivity to the informational state of conspecifics when performing social gazing behaviour. Aside from the “mind reading” hypothesis, three other hypotheses could however be put forward to explain why the gazing behaviour of the discoverer varied with the epistemic state of its conspecifics. The “saliency” hypothesis states that informed individuals perform more conspicuous behaviours than uninformed ones, and thus attract the attention of the discoverer. The “behaviour reading” hypothesis supposes that the discoverer strategically adjusts its gazing behaviour based on the behaviour of other group members. In the neutral condition, informed individuals received more gazes than uninformed ones, despite the fact that the wood bark chips were not supposed to elicit conspicuous or odd behaviours from the L’Hoest’s monkeys. In light of this result, we discarded the “saliency” and the “behaviour-reading” hypotheses.

The “proximity” hypothesis states that discoverers looked more at the closest individuals, which had checked the content of the box themselves. Our results show that the closest individual was more likely gazed at if there were at least three individuals in the vicinity of the discoverer. Since the discoverer was most of the time surrounded by more than two individuals, our results suggest that individuals in social proximity attracted most of the social gazes. Thus, the “proximity” hypothesis cannot clearly be discarded. One may note that the “proximity” hypothesis could account for the social gazing behaviour of the L’Hoest’s monkeys only if informed individuals remained in close proximity with the discoverer during the experiment, which could not be tested with our data. In sum, we could safely discard both “saliency” and “behaviour-reading” hypotheses, but it remained unclear whether the “proximity” hypothesis could be discarded.

Our results suggest that L’Hoest’s monkeys might be sensitive to the informational state of conspecifics when performing social gazing behaviour. However, this depended on the nature of the stimulus, as we did not find differences in the gazing behaviour in the aversive condition.

In chimpanzees, adjustment of vocal communication according to audience’s knowledge states has been found in alarm contexts (Crockford et al. 2012, 2017). Similarly to vocalizations, social gazes may qualify as communicative signals (i.e., looking at others to attract their attention to an item in the environment). Our results suggest that L’Hoest’s monkeys do not use social gazes to preferentially warn uninformed conspecifics. Besides, the fact that informed individuals were preferentially gazed at when the content of the box was food (i.e. mealworms) could be interpreted as assessment of the behaviour of knowledgeable competitors to best prepare to the release of food, as revealed in mangabeys (*Cercocebus torquatus*, Blois-Heulin and Girona 1999). Interestingly, in some experiments where a subject was released in a room with a dominant competitor, the subject behaved differently according to whether the competitor had seen a hidden food or not, suggesting that they understand that seeing is knowing (*Pan troglodytes*, Hare et al. 2001; *Cebus apella*, Hare et al. 2003). In our experiment, the discoverer is simultaneously facing potential competitors who have seen the hidden and unreachable food and others who have not. This experiment is an even more realistic context fitting with the socio-ecology of the tested species, which lives in groups of 10 to 17 individuals.

Differences in social gaze pattern in the neutral condition suggest that L’Hoest’s monkeys might be able to track conspecifics’ knowledge and adjust their social gazing behaviour accordingly (“mind reading” hypothesis). Future studies in L’Hoest’s monkeys or other animal species should yet elaborate on our protocol to disentangle the underlying mechanisms involved in the social gazing behaviour.

Overall, our study calls for a more systematic assessment of the “theory of mind” capacities beyond ape species. Investigating the diversity of mind-reading abilities of primates can unveil sophisticated cognitive capacities that were thought to be uniquely human until recently, and help to understand how they emerged from primitive systems (Martin and Santos 2016; Meunier 2017).

## Acknowledgments

The Authors thank the caregivers of La Menagerie (Christelle Hano, Christophe Bazin, Thibaud Pers, Tommy Tanchoux and Valérie Martinez) and the Director (Michel Saint Jalme) for administrative and technical support, and for approving the study. We are also grateful to Christof Neumann, Philippe Schlenker, Emmanuel Chemla and Pierre Jacob for conceptual and statistical support, and to Thomas T. Struhsaker for helpful information on the behaviour of L’Hoest’s monkeys.

## Declarations

### Ethics approval

All procedures were in accordance with the ethical standards of the institution at which the study was conducted.

### Consent to participate

Not applicable.

### Consent for publication

Not applicable.

### Availability of data and material

All the data and material used in this study are available as supplementary material.

### Code availability

Code used for statistical analysis is available as supplementary material

### Authors contribution

GD, LR and AM designed the study. LR collected the data. LR, AM, GD and MB performed the statistical analyses. All authors drafted the article and approved the final version of the manuscript.

